## Supplemental figures for "Targeting IRE1α improves insulin sensitivity and thermogenesis and suppresses metabolically active adipose tissue macrophages in male obese mice"

### Slide 1
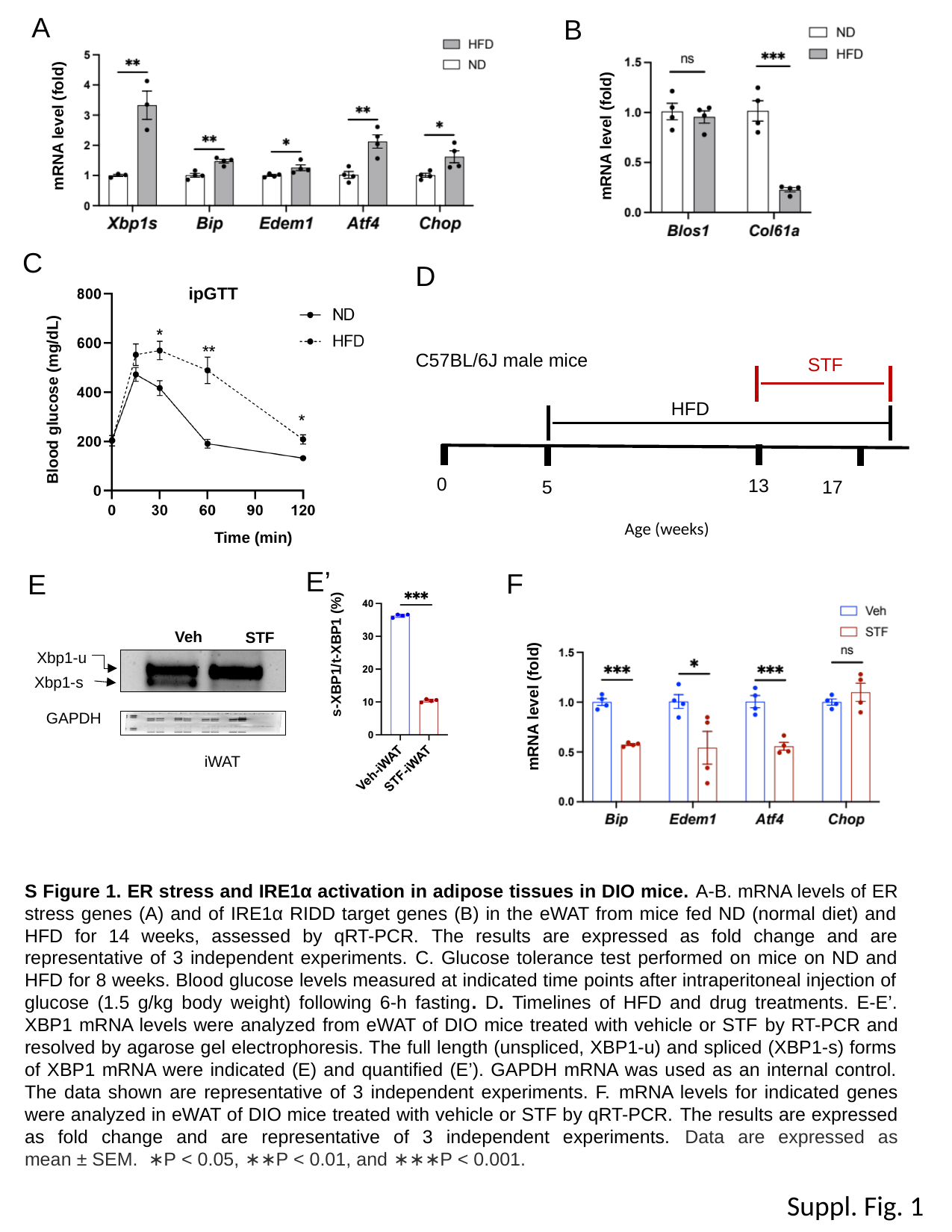

A
B
mRNA level (fold)
mRNA level (fold)
C
D
ipGTT
Blood glucose (mg/dL)
Time (min)
C57BL/6J male mice
STF
HFD
 0
 13
 17
 5
Age (weeks)
s-XBP1/t-XBP1 (%)
E’
F
E
Veh
STF
iWAT
Xbp1-u
Xbp1-s
GAPDH
mRNA level (fold)
S Figure 1. ER stress and IRE1α activation in adipose tissues in DIO mice. A-B. mRNA levels of ER stress genes (A) and of IRE1α RIDD target genes (B) in the eWAT from mice fed ND (normal diet) and HFD for 14 weeks, assessed by qRT-PCR. The results are expressed as fold change and are representative of 3 independent experiments. C. Glucose tolerance test performed on mice on ND and HFD for 8 weeks. Blood glucose levels measured at indicated time points after intraperitoneal injection of glucose (1.5 g/kg body weight) following 6-h fasting. D. Timelines of HFD and drug treatments. E-E’. XBP1 mRNA levels were analyzed from eWAT of DIO mice treated with vehicle or STF by RT-PCR and resolved by agarose gel electrophoresis. The full length (unspliced, XBP1-u) and spliced (XBP1-s) forms of XBP1 mRNA were indicated (E) and quantified (E’). GAPDH mRNA was used as an internal control. The data shown are representative of 3 independent experiments. F. mRNA levels for indicated genes were analyzed in eWAT of DIO mice treated with vehicle or STF by qRT-PCR. The results are expressed as fold change and are representative of 3 independent experiments. Data are expressed as mean ± SEM.  ∗P < 0.05, ∗∗P < 0.01, and ∗∗∗P < 0.001.
Suppl. Fig. 1

### Slide 2
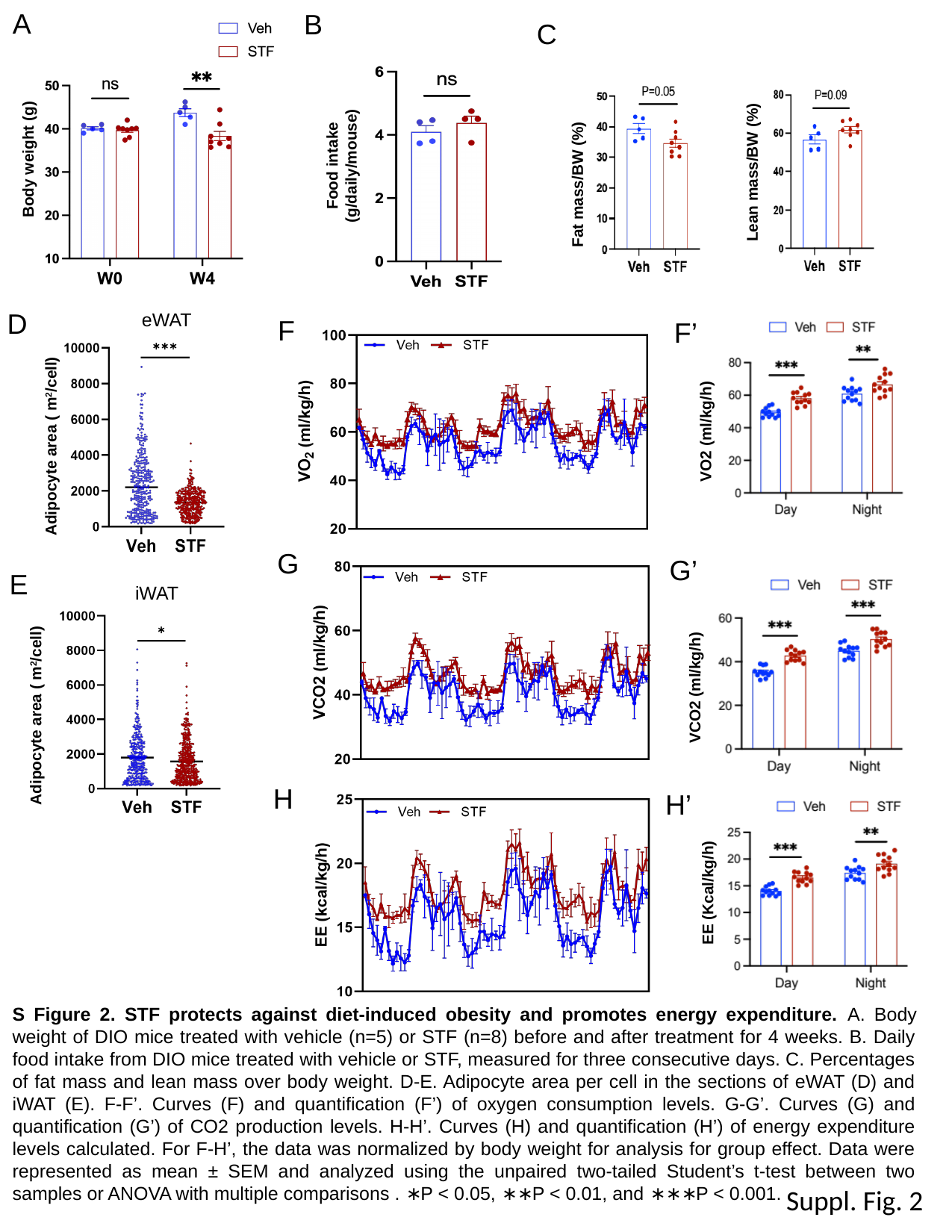

A
B
C
Lean mass/BW (%)
Fat mass/BW (%)
Food intake (g/daily/mouse)
Body weight (g)
D
eWAT
F
F’
VO2 (ml/kg/h)
G
G’
E
iWAT
VCO2 (ml/kg/h)
H
H’
EE (Kcal/kg/h)
S Figure 2. STF protects against diet-induced obesity and promotes energy expenditure. A. Body weight of DIO mice treated with vehicle (n=5) or STF (n=8) before and after treatment for 4 weeks. B. Daily food intake from DIO mice treated with vehicle or STF, measured for three consecutive days. C. Percentages of fat mass and lean mass over body weight. D-E. Adipocyte area per cell in the sections of eWAT (D) and iWAT (E). F-F’. Curves (F) and quantification (F’) of oxygen consumption levels. G-G’. Curves (G) and quantification (G’) of CO2 production levels. H-H’. Curves (H) and quantification (H’) of energy expenditure levels calculated. For F-H’, the data was normalized by body weight for analysis for group effect. Data were represented as mean ± SEM and analyzed using the unpaired two-tailed Student’s t-test between two samples or ANOVA with multiple comparisons . ∗P < 0.05, ∗∗P < 0.01, and ∗∗∗P < 0.001.
Suppl. Fig. 2

### Slide 3
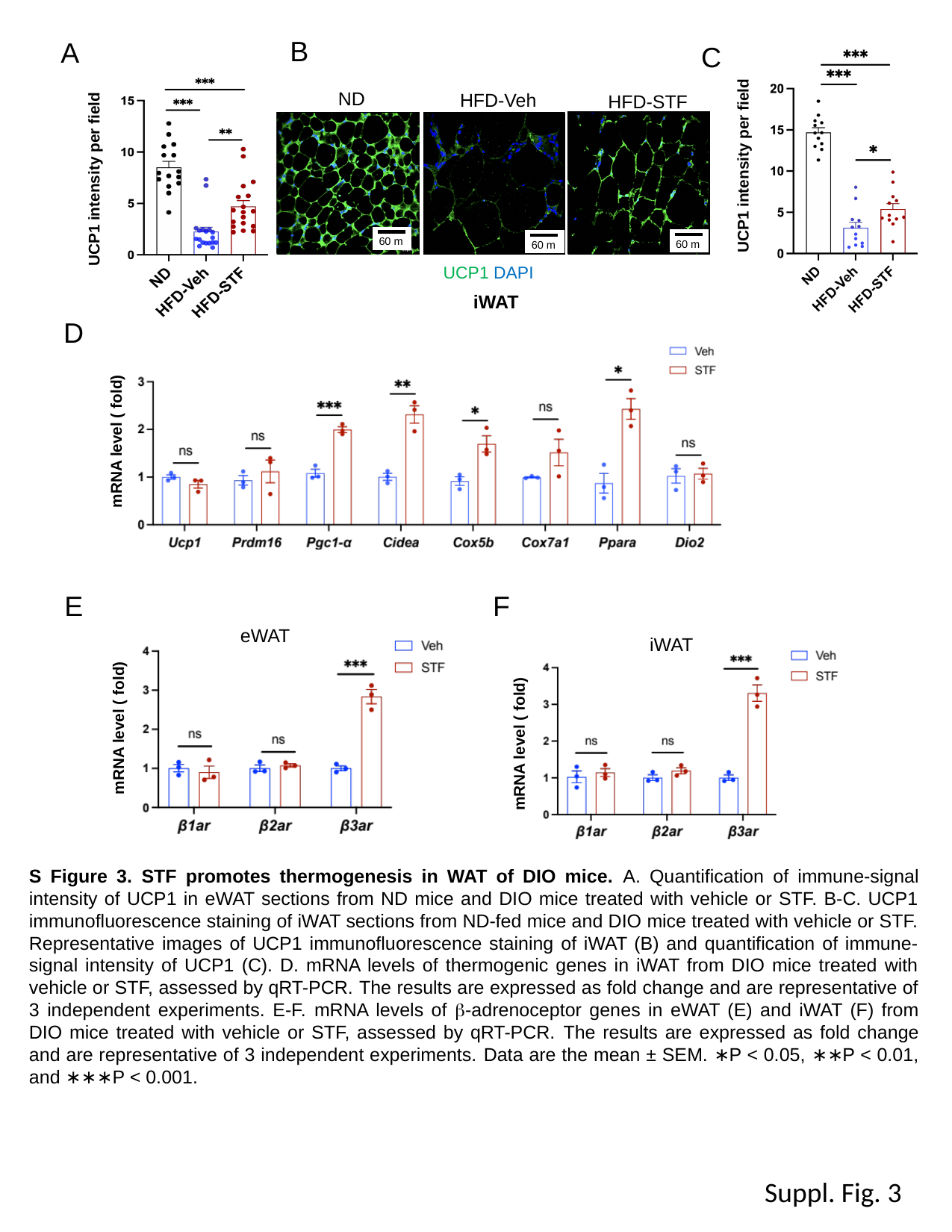

B
A
C
UCP1 intensity per field
ND
HFD-Veh
HFD-STF
iWAT
UCP1 intensity per field
UCP1 DAPI
D
mRNA level ( fold)
F
E
eWAT
mRNA level ( fold)
iWAT
mRNA level ( fold)
S Figure 3. STF promotes thermogenesis in WAT of DIO mice. A. Quantification of immune-signal intensity of UCP1 in eWAT sections from ND mice and DIO mice treated with vehicle or STF. B-C. UCP1 immunofluorescence staining of iWAT sections from ND-fed mice and DIO mice treated with vehicle or STF. Representative images of UCP1 immunofluorescence staining of iWAT (B) and quantification of immune-signal intensity of UCP1 (C). D. mRNA levels of thermogenic genes in iWAT from DIO mice treated with vehicle or STF, assessed by qRT-PCR. The results are expressed as fold change and are representative of 3 independent experiments. E-F. mRNA levels of β-adrenoceptor genes in eWAT (E) and iWAT (F) from DIO mice treated with vehicle or STF, assessed by qRT-PCR. The results are expressed as fold change and are representative of 3 independent experiments. Data are the mean ± SEM. ∗P < 0.05, ∗∗P < 0.01, and ∗∗∗P < 0.001.
Suppl. Fig. 3

### Slide 4
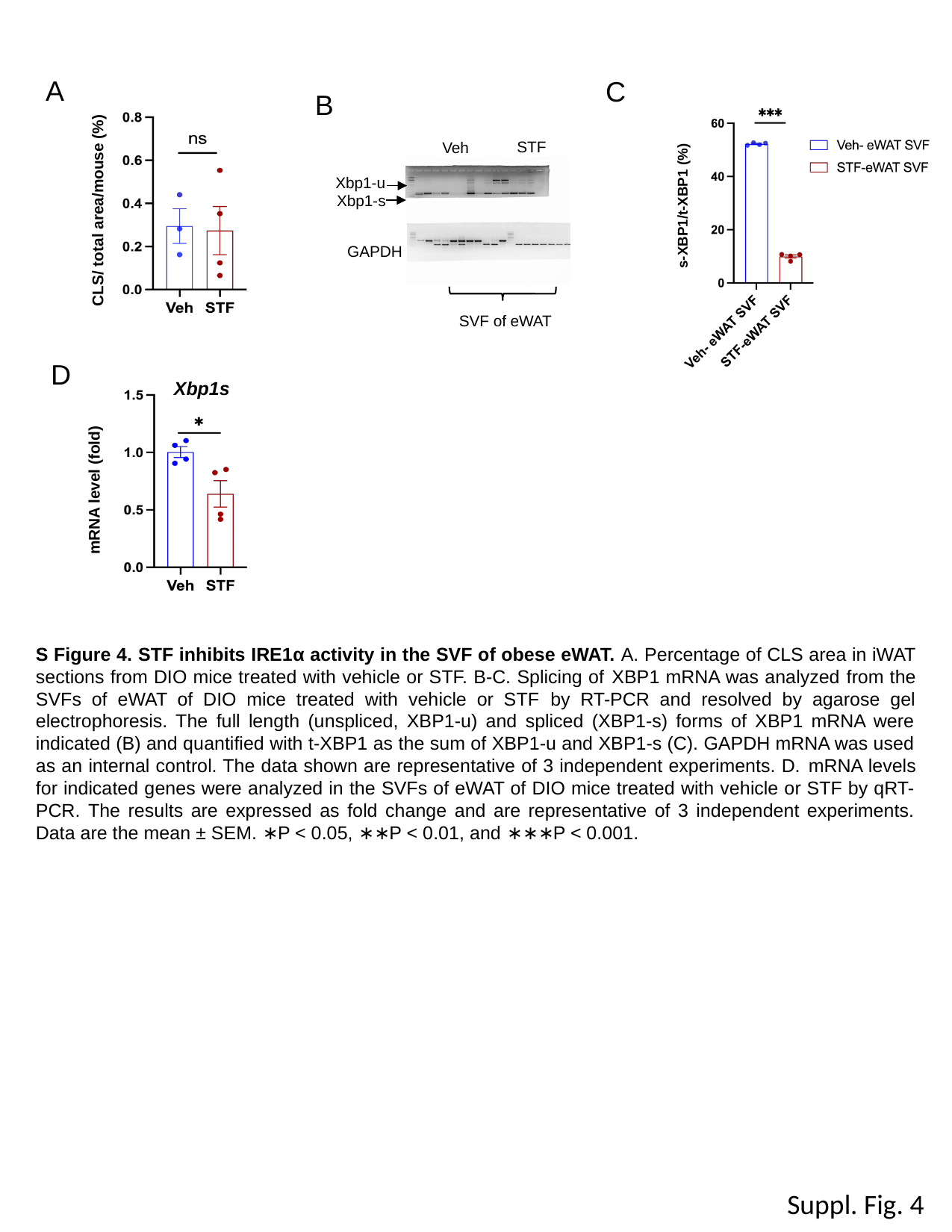

A
C
B
s-XBP1/t-XBP1 (%)
STF
Veh
Xbp1-u
Xbp1-s
GAPDH
SVF of eWAT
CLS/ total area/mouse (%)
D
Xbp1s
mRNA level (fold)
S Figure 4. STF inhibits IRE1α activity in the SVF of obese eWAT. A. Percentage of CLS area in iWAT sections from DIO mice treated with vehicle or STF. B-C. Splicing of XBP1 mRNA was analyzed from the SVFs of eWAT of DIO mice treated with vehicle or STF by RT-PCR and resolved by agarose gel electrophoresis. The full length (unspliced, XBP1-u) and spliced (XBP1-s) forms of XBP1 mRNA were indicated (B) and quantified with t-XBP1 as the sum of XBP1-u and XBP1-s (C). GAPDH mRNA was used as an internal control. The data shown are representative of 3 independent experiments. D. mRNA levels for indicated genes were analyzed in the SVFs of eWAT of DIO mice treated with vehicle or STF by qRT-PCR. The results are expressed as fold change and are representative of 3 independent experiments. Data are the mean ± SEM. ∗P < 0.05, ∗∗P < 0.01, and ∗∗∗P < 0.001.
Suppl. Fig. 4

### Slide 5
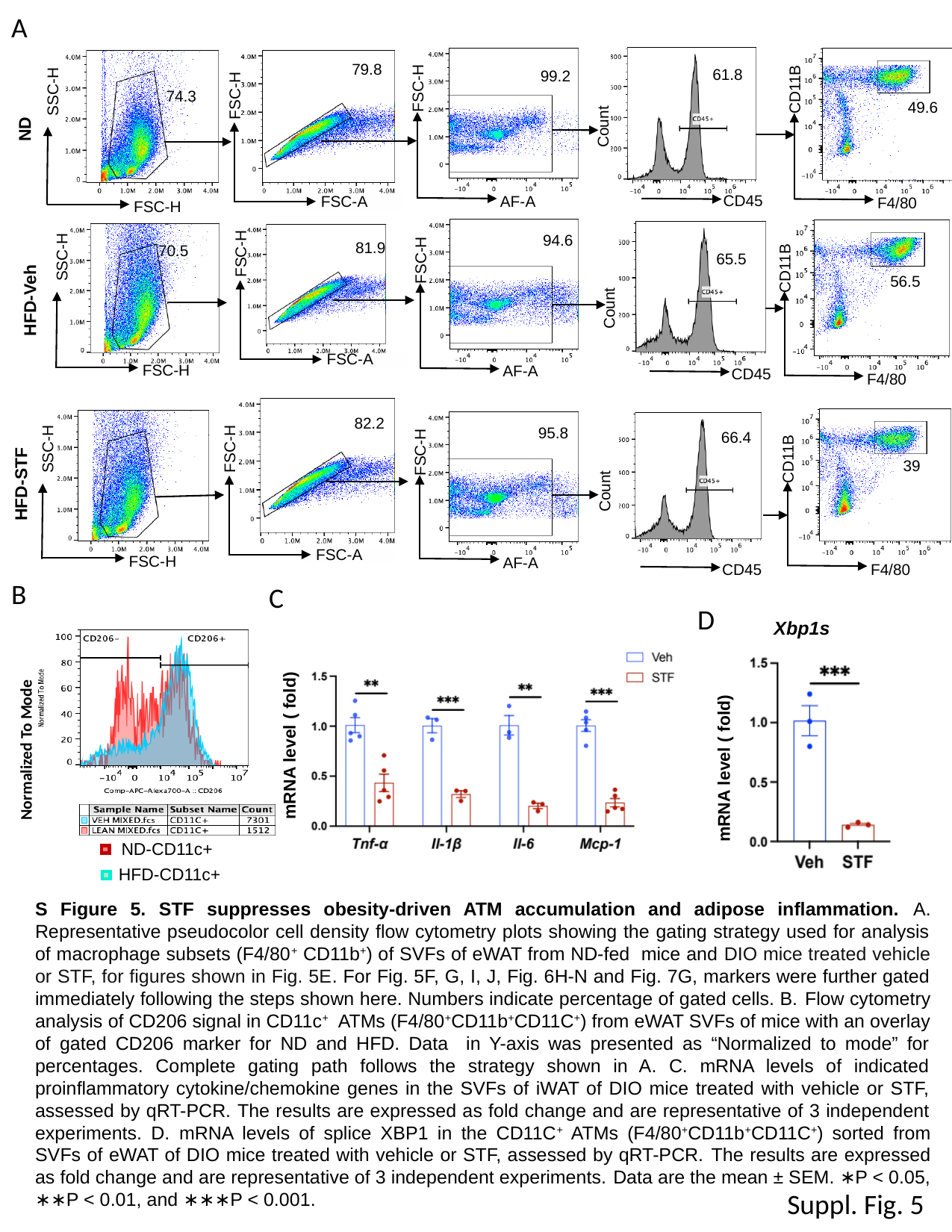

A
CD11B
F4/80
49.6
FSC-H
AF-A
79.8
SSC-H
FSC-H
61.8
FSC-H
FSC-A
99.2
74.3
ND
Count
CD45
FSC-H
FSC-A
SSC-H
FSC-H
FSC-H
AF-A
94.6
CD11B
F4/80
81.9
70.5
65.5
56.5
HFD-Veh
Count
CD45
68.7
82.2
SSC-H
FSC-H
FSC-H
FSC-A
FSC-H
AF-A
95.8
66.4
CD11B
F4/80
HFD-STF
39
Count
CD45
B
C
D
Xbp1s
mRNA level ( fold)
mRNA level ( fold)
Normalized To Mode
ND-CD11c+
HFD-CD11c+
S Figure 5. STF suppresses obesity-driven ATM accumulation and adipose inflammation. A. Representative pseudocolor cell density flow cytometry plots showing the gating strategy used for analysis of macrophage subsets (F4/80^+^ CD11b^+^) of SVFs of eWAT from ND-fed mice and DIO mice treated vehicle or STF, for figures shown in Fig. 5E. For Fig. 5F, G, I, J, Fig. 6H-N and Fig. 7G, markers were further gated immediately following the steps shown here. Numbers indicate percentage of gated cells. B. Flow cytometry analysis of CD206 signal in CD11c^+^ ATMs (F4/80^+^CD11b^+^CD11C^+^) from eWAT SVFs of mice with an overlay of gated CD206 marker for ND and HFD. Data in Y-axis was presented as “Normalized to mode” for percentages. Complete gating path follows the strategy shown in A. C. mRNA levels of indicated proinflammatory cytokine/chemokine genes in the SVFs of iWAT of DIO mice treated with vehicle or STF, assessed by qRT-PCR. The results are expressed as fold change and are representative of 3 independent experiments. D. mRNA levels of splice XBP1 in the CD11C^+^ ATMs (F4/80^+^CD11b^+^CD11C^+^) sorted from SVFs of eWAT of DIO mice treated with vehicle or STF, assessed by qRT-PCR. The results are expressed as fold change and are representative of 3 independent experiments. Data are the mean ± SEM. ∗P < 0.05, ∗∗P < 0.01, and ∗∗∗P < 0.001.
Suppl. Fig. 5

### Slide 6
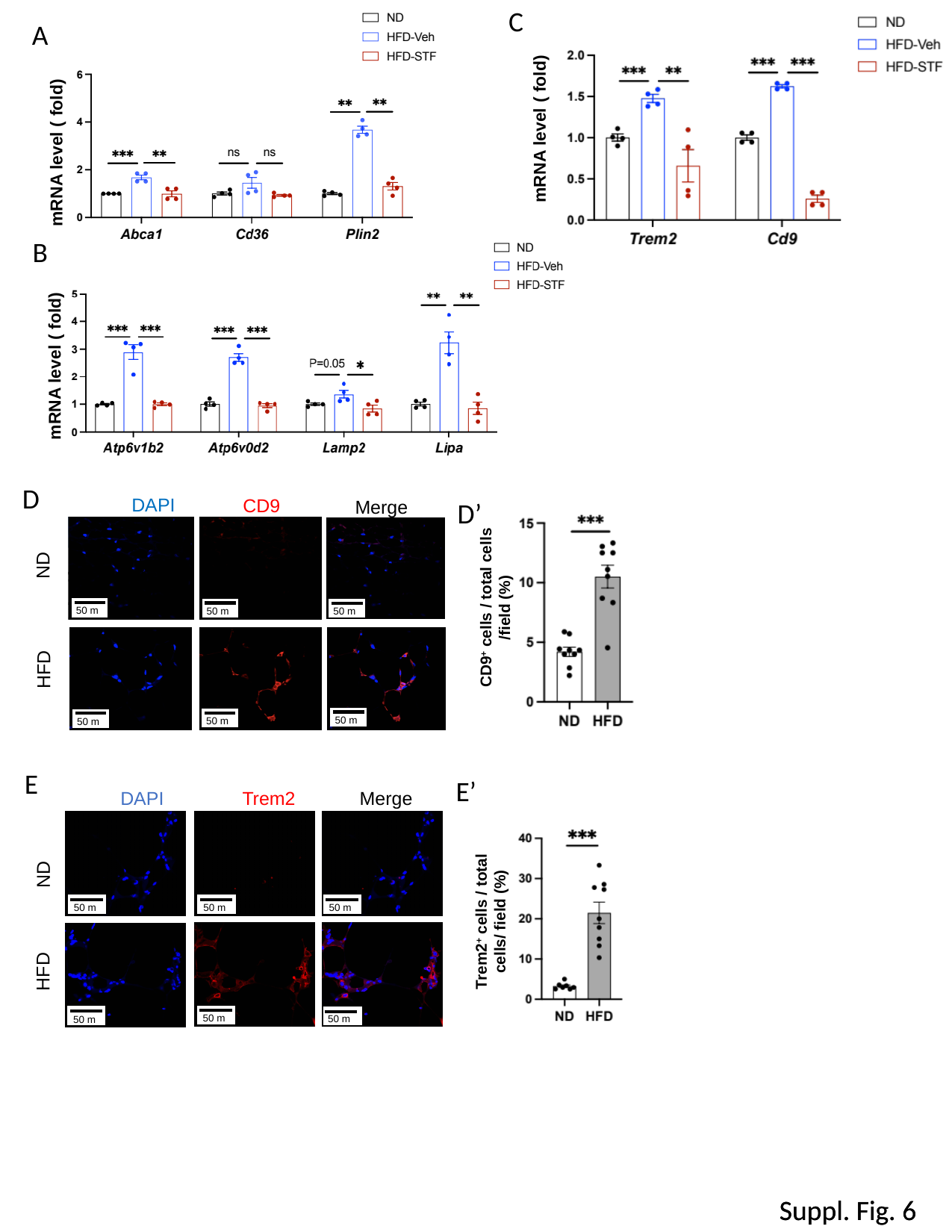

C
A
mRNA level ( fold)
mRNA level ( fold)
B
mRNA level ( fold)
D
DAPI
CD9
Merge
ND
HFD
D’
CD9^+^ cells / total cells /field (%)
E
E’
DAPI
Trem2
Merge
ND
HFD
Trem2^+^ cells / total cells/ field (%)
Suppl. Fig. 6
Suppl. Fig. 6

### Slide 7
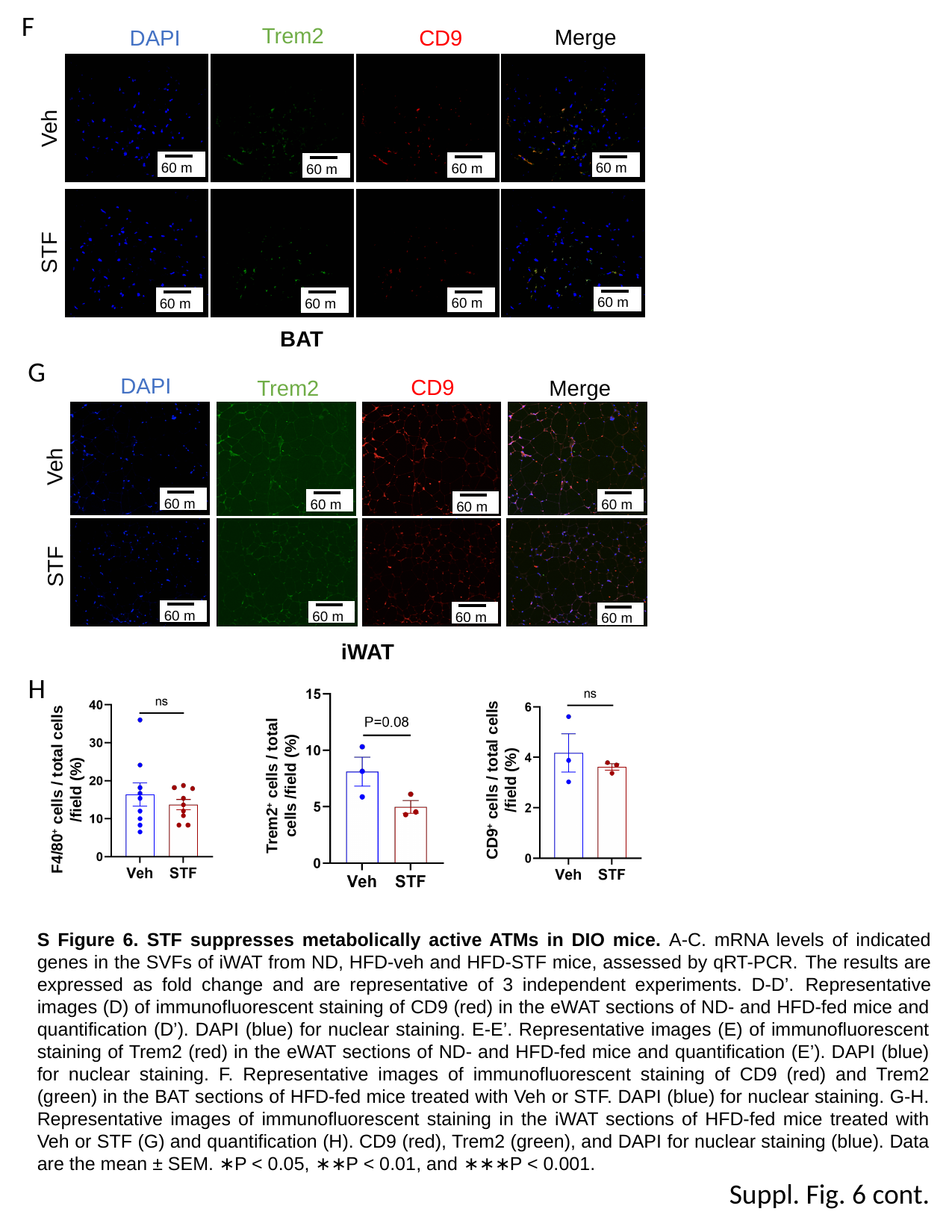

F
Trem2
Merge
DAPI
CD9
Veh
STF
BAT
G
DAPI
CD9
Merge
Trem2
Veh
STF
iWAT
H
CD9^+^ cells / total cells /field (%)
Trem2^+^ cells / total cells /field (%)
F4/80^+^ cells / total cells /field (%)
S Figure 6. STF suppresses metabolically active ATMs in DIO mice. A-C. mRNA levels of indicated genes in the SVFs of iWAT from ND, HFD-veh and HFD-STF mice, assessed by qRT-PCR. The results are expressed as fold change and are representative of 3 independent experiments. D-D’. Representative images (D) of immunofluorescent staining of CD9 (red) in the eWAT sections of ND- and HFD-fed mice and quantification (D’). DAPI (blue) for nuclear staining. E-E’. Representative images (E) of immunofluorescent staining of Trem2 (red) in the eWAT sections of ND- and HFD-fed mice and quantification (E’). DAPI (blue) for nuclear staining. F. Representative images of immunofluorescent staining of CD9 (red) and Trem2 (green) in the BAT sections of HFD-fed mice treated with Veh or STF. DAPI (blue) for nuclear staining. G-H. Representative images of immunofluorescent staining in the iWAT sections of HFD-fed mice treated with Veh or STF (G) and quantification (H). CD9 (red), Trem2 (green), and DAPI for nuclear staining (blue). Data are the mean ± SEM. ∗P < 0.05, ∗∗P < 0.01, and ∗∗∗P < 0.001.
Suppl. Fig. 6 cont.

### Slide 8
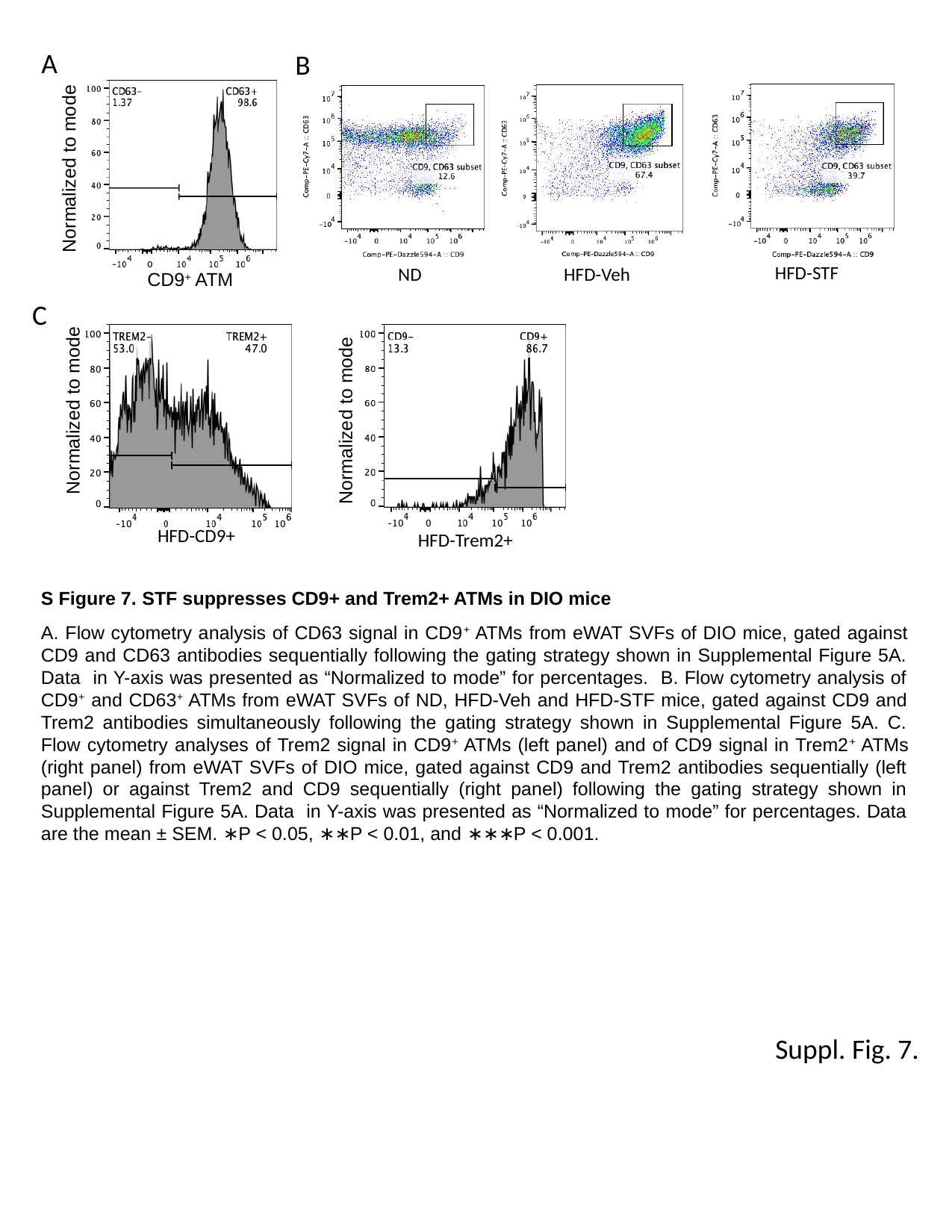

A
CD9^+^ ATM
B
HFD-STF
HFD-Veh
ND
Normalized to mode
C
Normalized to mode
HFD-CD9+
Normalized to mode
HFD-Trem2+
S Figure 7. STF suppresses CD9+ and Trem2+ ATMs in DIO mice
A. Flow cytometry analysis of CD63 signal in CD9^+^ ATMs from eWAT SVFs of DIO mice, gated against CD9 and CD63 antibodies sequentially following the gating strategy shown in Supplemental Figure 5A. Data in Y-axis was presented as “Normalized to mode” for percentages. B. Flow cytometry analysis of CD9^+^ and CD63^+^ ATMs from eWAT SVFs of ND, HFD-Veh and HFD-STF mice, gated against CD9 and Trem2 antibodies simultaneously following the gating strategy shown in Supplemental Figure 5A. C. Flow cytometry analyses of Trem2 signal in CD9^+^ ATMs (left panel) and of CD9 signal in Trem2^+^ ATMs (right panel) from eWAT SVFs of DIO mice, gated against CD9 and Trem2 antibodies sequentially (left panel) or against Trem2 and CD9 sequentially (right panel) following the gating strategy shown in Supplemental Figure 5A. Data in Y-axis was presented as “Normalized to mode” for percentages. Data are the mean ± SEM. ∗P < 0.05, ∗∗P < 0.01, and ∗∗∗P < 0.001.
Suppl. Fig. 7.

### Slide 9
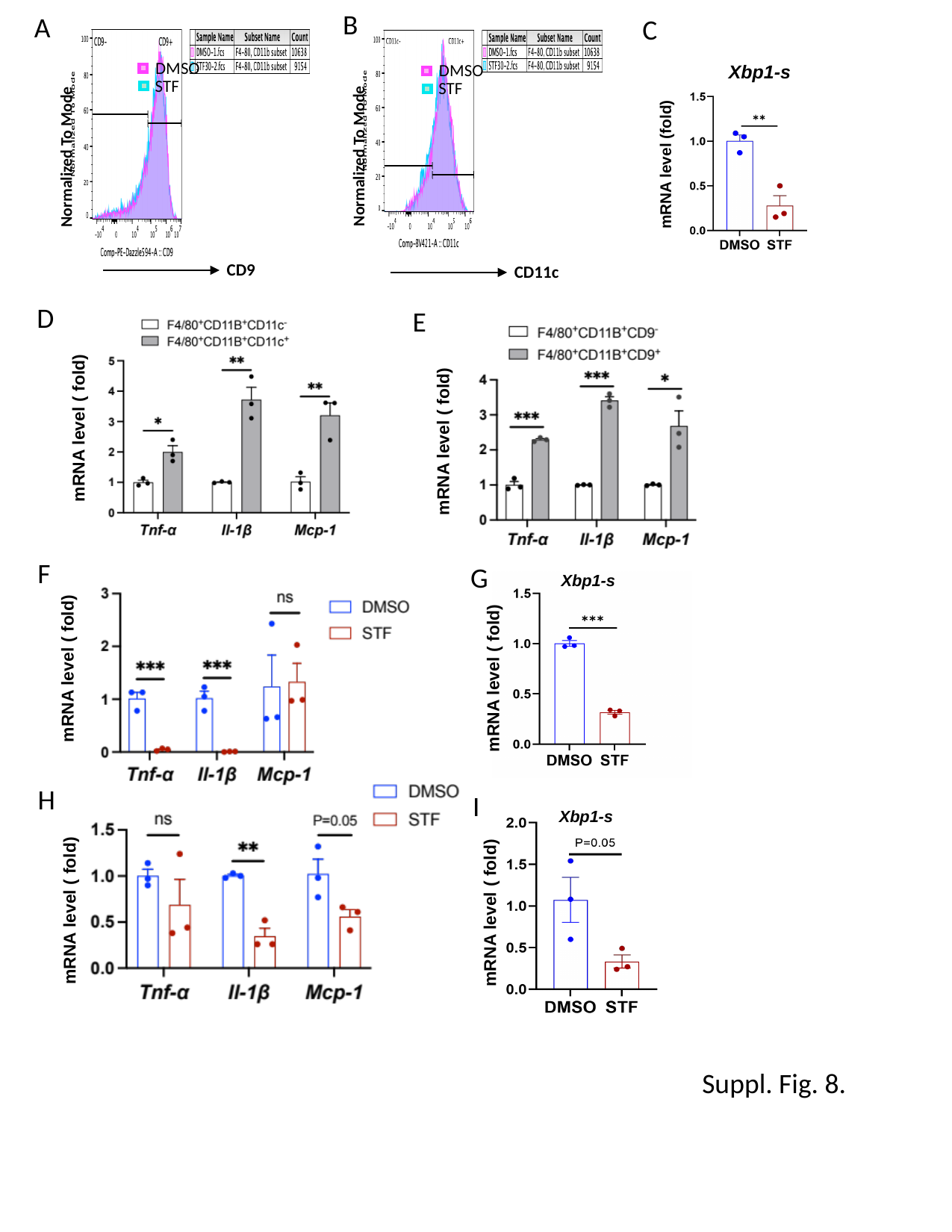

B
A
C
CD9
DMSO
STF
Normalized To Mode
CD11c
DMSO
STF
Normalized To Mode
Xbp1-s
mRNA level (fold)
D
E
mRNA level ( fold)
mRNA level ( fold)
F
G
Xbp1-s
mRNA level ( fold)
mRNA level ( fold)
H
I
Xbp1-s
mRNA level ( fold)
mRNA level ( fold)
Suppl. Fig. 8.

### Slide 10
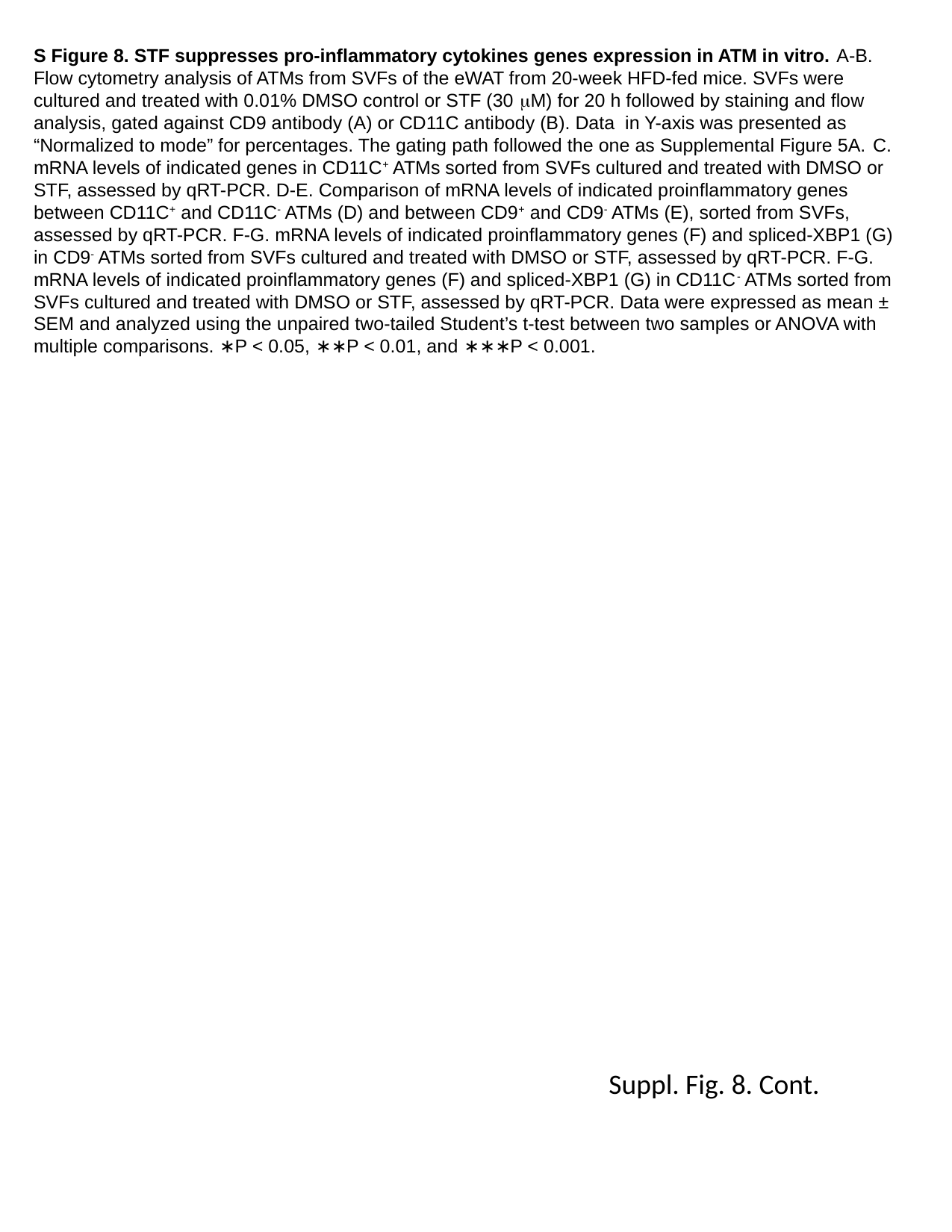

S Figure 8. STF suppresses pro-inflammatory cytokines genes expression in ATM in vitro. A-B. Flow cytometry analysis of ATMs from SVFs of the eWAT from 20-week HFD-fed mice. SVFs were cultured and treated with 0.01% DMSO control or STF (30 μM) for 20 h followed by staining and flow analysis, gated against CD9 antibody (A) or CD11C antibody (B). Data in Y-axis was presented as “Normalized to mode” for percentages. The gating path followed the one as Supplemental Figure 5A. C. mRNA levels of indicated genes in CD11C^+^ ATMs sorted from SVFs cultured and treated with DMSO or STF, assessed by qRT-PCR. D-E. Comparison of mRNA levels of indicated proinflammatory genes between CD11C^+^ and CD11C^-^ ATMs (D) and between CD9^+^ and CD9^-^ ATMs (E), sorted from SVFs, assessed by qRT-PCR. F-G. mRNA levels of indicated proinflammatory genes (F) and spliced-XBP1 (G) in CD9^-^ ATMs sorted from SVFs cultured and treated with DMSO or STF, assessed by qRT-PCR. F-G. mRNA levels of indicated proinflammatory genes (F) and spliced-XBP1 (G) in CD11C^-^ ATMs sorted from SVFs cultured and treated with DMSO or STF, assessed by qRT-PCR. Data were expressed as mean ± SEM and analyzed using the unpaired two-tailed Student’s t-test between two samples or ANOVA with multiple comparisons. ∗P < 0.05, ∗∗P < 0.01, and ∗∗∗P < 0.001.
Suppl. Fig. 8. Cont.

### Slide 11
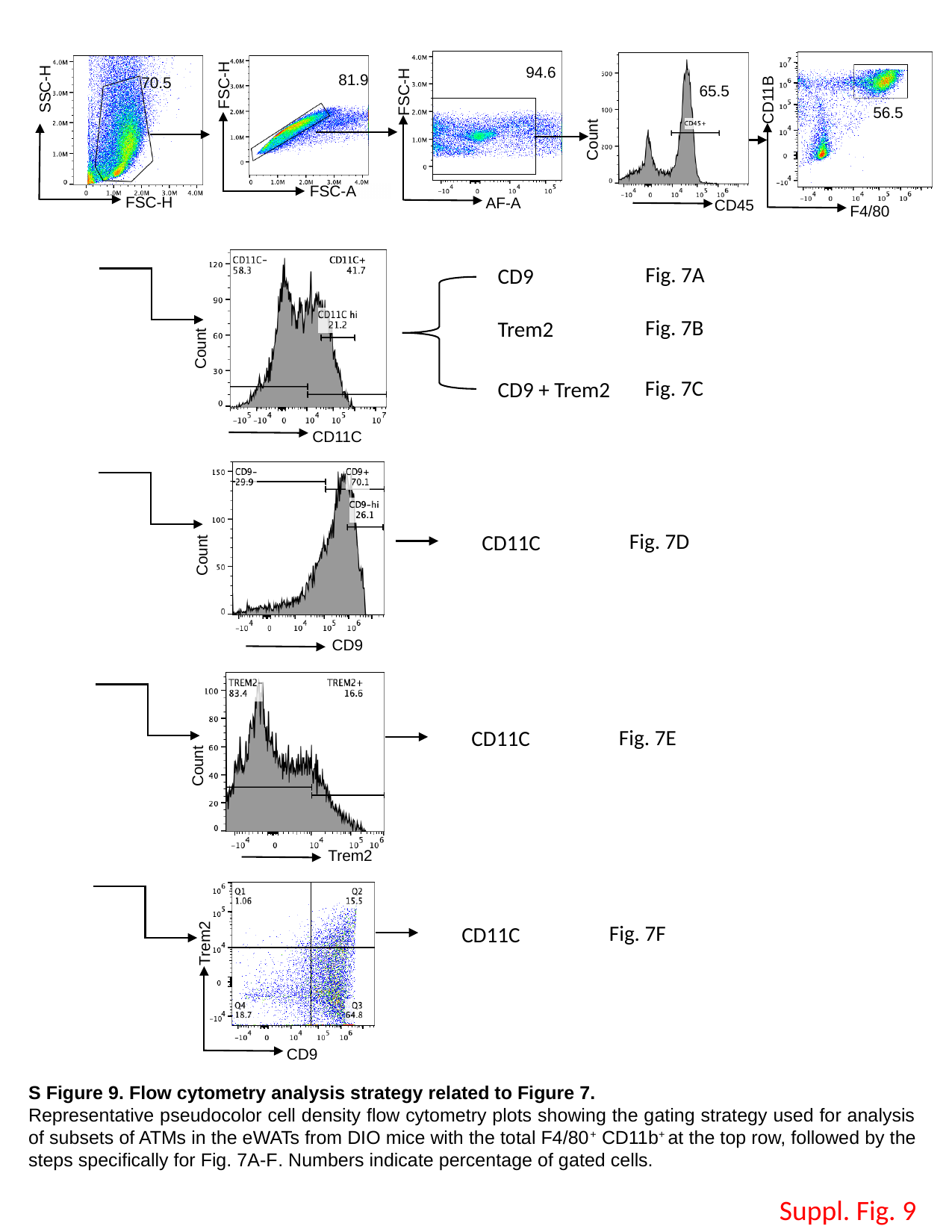

FSC-H
FSC-A
SSC-H
FSC-H
FSC-H
AF-A
94.6
CD11B
F4/80
81.9
70.5
65.5
56.5
Count
CD45
Fig. 7A
CD9
Fig. 7B
Trem2
Count
CD11C
Fig. 7C
CD9 + Trem2
Fig. 7D
Count
CD9
CD11C
Fig. 7E
CD11C
Count
Trem2
Trem2
CD9
Fig. 7F
CD11C
S Figure 9. Flow cytometry analysis strategy related to Figure 7.
Representative pseudocolor cell density flow cytometry plots showing the gating strategy used for analysis of subsets of ATMs in the eWATs from DIO mice with the total F4/80^+^ CD11b^+^ at the top row, followed by the steps specifically for Fig. 7A-F. Numbers indicate percentage of gated cells.
Suppl. Fig. 9

### Slide 12
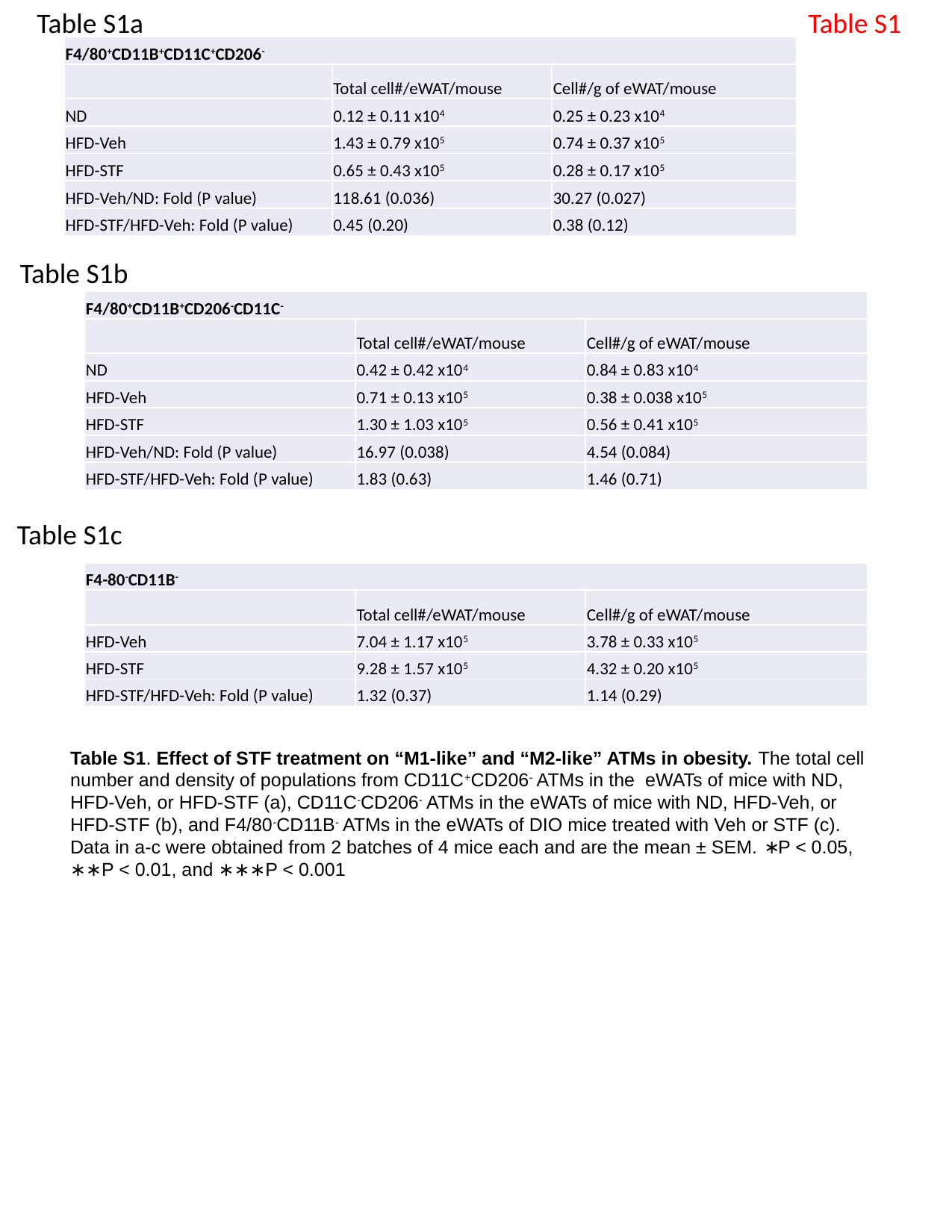

Table S1a
Table S1
| F4/80^+^CD11B^+^CD11C^+^CD206^-^ | | |
| --- | --- | --- |
| | Total cell#/eWAT/mouse | Cell#/g of eWAT/mouse |
| ND | 0.12 ± 0.11 x10^4^ | 0.25 ± 0.23 x10^4^ |
| HFD-Veh | 1.43 ± 0.79 x10^5^ | 0.74 ± 0.37 x10^5^ |
| HFD-STF | 0.65 ± 0.43 x10^5^ | 0.28 ± 0.17 x10^5^ |
| HFD-Veh/ND: Fold (P value) | 118.61 (0.036) | 30.27 (0.027) |
| HFD-STF/HFD-Veh: Fold (P value) | 0.45 (0.20) | 0.38 (0.12) |
Table S1b
| F4/80^+^CD11B^+^CD206^-^CD11C^-^ | | |
| --- | --- | --- |
| | Total cell#/eWAT/mouse | Cell#/g of eWAT/mouse |
| ND | 0.42 ± 0.42 x10^4^ | 0.84 ± 0.83 x10^4^ |
| HFD-Veh | 0.71 ± 0.13 x10^5^ | 0.38 ± 0.038 x10^5^ |
| HFD-STF | 1.30 ± 1.03 x10^5^ | 0.56 ± 0.41 x10^5^ |
| HFD-Veh/ND: Fold (P value) | 16.97 (0.038) | 4.54 (0.084) |
| HFD-STF/HFD-Veh: Fold (P value) | 1.83 (0.63) | 1.46 (0.71) |
Table S1c
| F4-80^-^CD11B^-^ | | |
| --- | --- | --- |
| | Total cell#/eWAT/mouse | Cell#/g of eWAT/mouse |
| HFD-Veh | 7.04 ± 1.17 x10^5^ | 3.78 ± 0.33 x10^5^ |
| HFD-STF | 9.28 ± 1.57 x10^5^ | 4.32 ± 0.20 x10^5^ |
| HFD-STF/HFD-Veh: Fold (P value) | 1.32 (0.37) | 1.14 (0.29) |
Table S1. Effect of STF treatment on “M1-like” and “M2-like” ATMs in obesity. The total cell number and density of populations from CD11C^+^CD206^-^ ATMs in the eWATs of mice with ND, HFD-Veh, or HFD-STF (a), CD11C^-^CD206^-^ ATMs in the eWATs of mice with ND, HFD-Veh, or HFD-STF (b), and F4/80^-^CD11B^-^ ATMs in the eWATs of DIO mice treated with Veh or STF (c). Data in a-c were obtained from 2 batches of 4 mice each and are the mean ± SEM. ∗P < 0.05, ∗∗P < 0.01, and ∗∗∗P < 0.001

### Slide 13
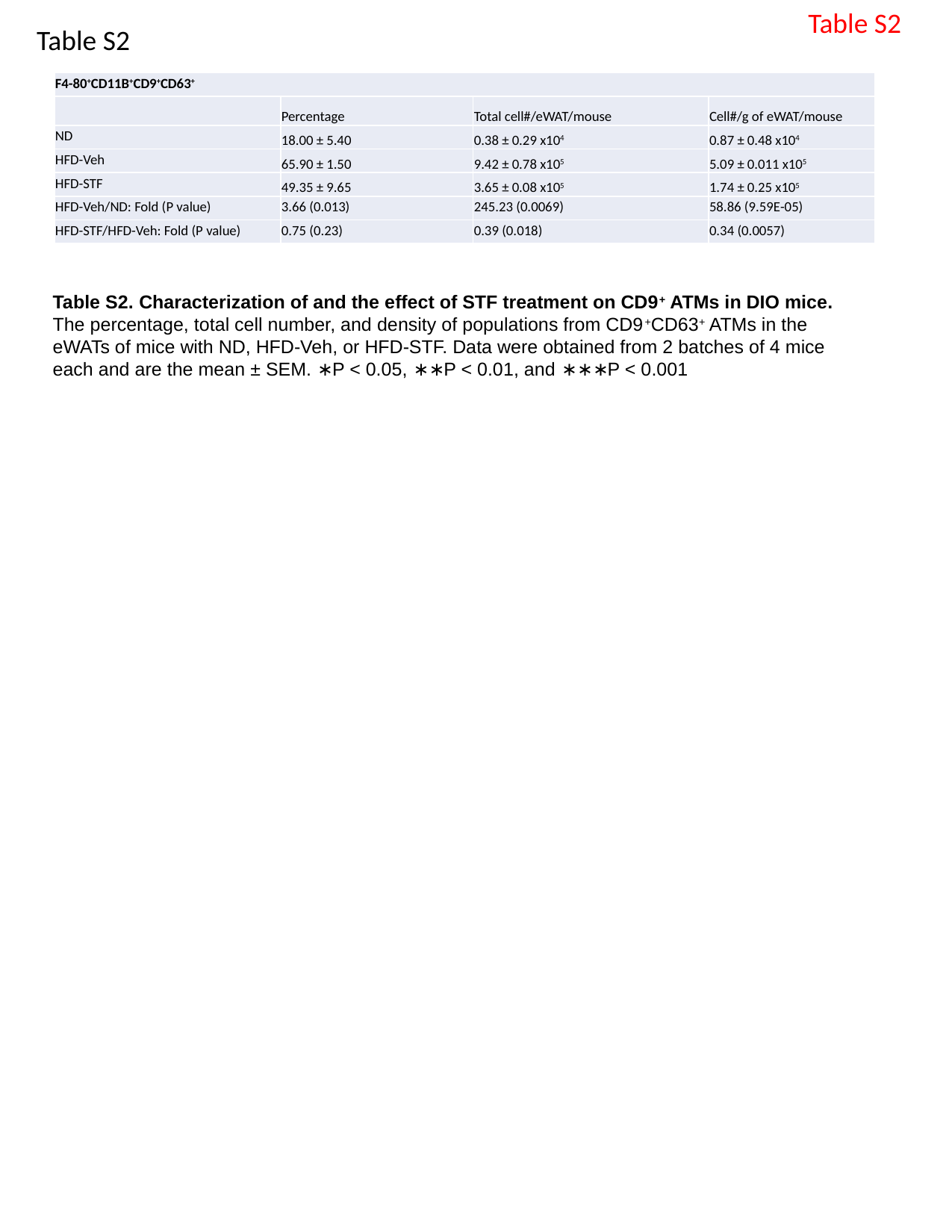

Table S2
Table S2
| F4-80^+^CD11B^+^CD9^+^CD63^+^ | | | |
| --- | --- | --- | --- |
| | Percentage | Total cell#/eWAT/mouse | Cell#/g of eWAT/mouse |
| ND | 18.00 ± 5.40 | 0.38 ± 0.29 x10^4^ | 0.87 ± 0.48 x10^4^ |
| HFD-Veh | 65.90 ± 1.50 | 9.42 ± 0.78 x10^5^ | 5.09 ± 0.011 x10^5^ |
| HFD-STF | 49.35 ± 9.65 | 3.65 ± 0.08 x10^5^ | 1.74 ± 0.25 x10^5^ |
| HFD-Veh/ND: Fold (P value) | 3.66 (0.013) | 245.23 (0.0069) | 58.86 (9.59E-05) |
| HFD-STF/HFD-Veh: Fold (P value) | 0.75 (0.23) | 0.39 (0.018) | 0.34 (0.0057) |
Table S2. Characterization of and the effect of STF treatment on CD9^+^ ATMs in DIO mice. The percentage, total cell number, and density of populations from CD9^+^CD63^+^ ATMs in the eWATs of mice with ND, HFD-Veh, or HFD-STF. Data were obtained from 2 batches of 4 mice each and are the mean ± SEM. ∗P < 0.05, ∗∗P < 0.01, and ∗∗∗P < 0.001
